# The impact of face masks on face-to-face neural tracking of speech: auditory and visual obstacles

**DOI:** 10.1101/2024.02.12.577414

**Authors:** M. Fantoni, A. Federici, I. Camponogara, G. Handjaras, A. Martinelli, E. Bednaya, E. Ricciardi, F. Pavani, D. Bottari

## Abstract

Face masks provide fundamental protection against the transmission of respiratory viruses but hamper communication. We estimated auditory and visual obstacles generated by face masks on communication by measuring the neural tracking of face-to-face speech. To this end, we recorded the EEG while participants were exposed to naturalistic audio-visual speech, embedded in multi-talker noise, in three contexts: (i) no-mask (audio-visual information was fully available), (ii) virtual mask (occluded lips, but intact audio), and (iii) real mask (occluded lips and degraded audio). The neural tracking of lip movements and the sound envelope of speech was measured through backward modeling, that is, by reconstructing stimulus properties from neural activity. Behaviorally, face masks increased listening -phonological-errors in speech content retrieval and perceived listening difficulty. At the neural level, we observed that the occlusion of the mouth abolished lip tracking and dampened neural tracking of the speech envelope at the earliest processing stages. Degraded acoustic information due to face mask filtering altered neural tracking at later processing stages instead. Finally, a consistent link emerged between the increment of listening perceived difficulty and the drop in reconstruction performance of speech envelope when attending to a speaker wearing a face mask. Results clearly dissociated the visual and auditory impacts of face masks on face-to-face neural tracking of speech. While face masks hampered the ability to predict and integrate audio-visual speech, the auditory filter generated by face masks impacted the neural processing stages typically associated with auditory selective attention. The link between perceived difficulty and neural tracking drop provided evidence of a major impact of face masks on the metacognitive levels subtending speech processing.

## 1. Introduction

The natural statistics of audio-visual speech are characterized by correlations and temporal correspondences between the visual changes in the mouth area and the sound envelope (Chandrasekaran et al., 2009). In turn, these signals are coupled to the neural processes of the listener, who exploits available information (Lakatos et al., 2019).

It was widely demonstrated that seeing the speaker’s facial movements, notably lip movements, enhances speech processing (Park et al., 2016). This boost is particularly relevant in challenging listening conditions, as in the case of noisy environments (Grant, Seitz, 2000; Ross et al., 2006) or hearing impairment (Song et al., 2014; Moradi et al., 2017). The benefit of multisensory input becomes fundamental to compensate for potential signal ambiguities (Holmes, 2009). When visual cues are lacking, such as when talking to a person wearing a face mask (like during the pandemic of COVID-19), comprehension becomes more challenging, requiring higher cognitive effort (Blackburn et al., 2019). Noteworthy is that face masks act both as a visual obstacle and an acoustic filter. They occlude the visibility of the mouth area and alter the sound emitted by the speaker (Corey et al., 2020; Haider et al., 2022). Thus, face masks represent a valuable natural context to estimate the relative contribution of visual and acoustic cues during audio-visual speech processing.

Behavioral studies have revealed that face masks impact speech intelligibility, especially when listening to speech in noise, whereas in quiet, there is limited interference (Brown et al., 2021). Different studies tested the degree to which various face masks impact speech comprehension and how this changes according to the background noise level (e.g., Brown et al., 2021; Toscano & Toscano, 2021). Moreover, it was documented that face masks affect the metacognitive dimensions of the listening experience. Indeed, it increases the perceived listening effort and reduces the confidence in what has been heard; also, it affects the meta-cognitive monitoring (i.e., the ability to correctly determine whether one’s performance was adequate or deficient) for speech recognition (Giovanelli et al., 2023). Thus, the presence of a face mask covering the lips of the speaker leads to reduced understanding, as well as other difficulties related to meta-cognitive assessments of the listening experience.

Changes at the brain processing level mirror these behavioral difficulties. Recent MEG investigation revealed that surgical face masks impact listeners’ neural tracking of acoustic features, such as sound envelope and spectral features (i.e., pitch and formant frequencies; Haider et al., 2022). Moreover, neural tracking of higher-level speech features such as phonemes and words’ onsets was found to be significantly hampered when listening to a speaker wearing a face mask in challenging auditory contexts (i.e., in the presence of additional speakers in the background). Nevertheless, these results could not be causally linked to the visual or acoustic filtering role exerted by the face mask, as the experimental design did not allow disambiguation between the two speech features.

Here, we aimed to precisely estimate visual and auditory obstacles generated by face masks by measuring the neural tracking of face-to-face speech. To this aim, we recorded the electroencephalography (EEG) while participants were exposed to naturalistic audio-visual speech stimuli in three conditions: (i) no-mask, (ii) virtual mask, and (iii) real mask. In the no-mask condition, visual and acoustic information was fully available to the listener. The acoustic information was preserved in the virtual mask condition, but a digital face mask hid the mouth area. Finally, the speaker’s lips were hidden in the real mask condition, and the acoustic information was degraded by the physical filter constituted by a surgical face mask. The auditory stream of the speaker was always embedded in babble noise (multiple talkers in the background) to promote audio-visual integration in all conditions. Indeed, since listening to speech in noise benefits from having concurrent audio-visual information, adding babble noise in the background could enhance the integration of auditory and visual speech signals (Golumbic, 2013; Crosse et al., 2015; Ahmed et al., 2023).

We measured participants’ accuracy at answering content questions and the relative committed errors, the confidence about their responses, and the perceived difficulty at attending speech in noise. Since behavior represents the outcome of many processes originating in the brain, we employed multivariate data analyses of the EEG to unveil the temporal dynamics subtending speech processing that could be unseen from performance alone. Neural tracking was estimated through backward modeling, aiming to reconstruct the visual or auditory input from neural data (Crosse et al., 2016; 2021). By measuring the neural tracking of lip movements and sound envelope across conditions, we assessed the perceptual cost of attending speech in noise when the speaker wore a face mask, distinguishing between the effects of the occlusion of the lips, degraded audio, and their combination. We predicted a better reconstruction of the lip movements and sound envelope in the no-mask condition, in which both the acoustic and visual cues were accessible, and a progressive dampening of neural tracking in the virtual and real mask conditions. We could selectively unravel the visual masking effect by contrasting the neural tracking of speech in the no-mask and virtual mask conditions (since these conditions share the same acoustic information). Instead, the comparison of the speech neural tracking in the virtual vs. real mask conditions allowed us to objectively disentangle the effect of the acoustic filter (present only in the real mask condition). Finally, we could estimate the combined effect of visual and acoustic obstacles by comparing the speech neural tracking between the no-mask and the real mask conditions. Results clearly revealed that the visual occlusion of the lips prevented lip tracking and hampered the neural tracking of the sound envelope in early processing stages, which benefited from the integration of audio-visual speech cues. Auditory filtering instead affected the neural tracking of the sound envelope at later stages, which are typically linked to higher-level auditory processing. Finally, the neural costs of listening to a speaker wearing a face mask were associated with behavioral costs measured at the metacognitive level (i.e., the perceived difficulty). Our results unequivocally revealed distinct effects of visual and auditory obstacles induced by face masks on the neural tracking of speech signals.

## 2. Materials and Methods

### 2.1 Participants

Thirty adults, all native Italian speakers, were enrolled in the study (N=30, Mean age=27.7 years, SD=2.1, Range: 21.3-31.3; 15 females). Participants did not report having a previous history of neurological disorders by self-report. The local Ethics Committee approved the study (protocol number: 1485/2017), which was conducted following the Declaration of Helsinki (2013). All participants signed informed consent before starting the experiment and received monetary compensation for their participation. We excluded one participant from the dataset due to the high number of interpolated channels (final sample EEG data: N=29 participants, mean age=27.6 years; 14 females). Due to technical issues, the behavioral data from one participant were not recorded (final sample behavioral data: N=28 participants, mean age=27.6 years; 13 females).

### 2.2 Stimuli

Speech stimuli consisted of 9 stories of approximately 3 minutes each. All stories were in Italian and chosen to be auto-conclusive. Stories were selected from the “Fiabe Italiane” by Italo Calvino, 1956 (“Italian tales”), “I Tacchini non ringraziano” by Andrea Camilleri (“Turkeys don’t give thanks”), and “Il momento è delicato” of Nicolò Ammaniti (“The delicate moment”).

A native Italian speaker read the stories while she was video recorded. Recordings were performed in a sound-attenuated chamber (BOXY, B-Beng s.r.l., Italy) at the IMT School for Advanced Study Lucca using an iPhone 7 (camera with 12MP, video resolution in HD, 720p with 30 fps, at a sampling frequency of 48000 kHz) and a condensation microphone (YC-LM10 II, Yichuang). The video frames comprised the speaker’s face with a homogeneous gray wall in the background. All stories were recorded twice, once while the speaker wore a surgical face mask (real mask condition) and once without it (no-mask condition). The no-mask recordings were also employed to create a third condition in which the speaker’s face was partially covered with a digital face mask artificially superimposed (virtual mask condition). The face mask used in the real mask condition was a Type IIR, three-layer, single-use medical face mask.

Each video began with silence, during which the speaker looked into the camera, then started reading the story’s title and began the storytelling. While reading, the speaker kept her eyes down (reading the text) and looked at the camera again only once the story was over. The speaker was asked to keep the head as still as possible. All audio-video recordings were imported in iMovie® (version 10.3.1; 1600 x 900 pixels resolution). To create the virtual mask condition for each video of the no-mask condition, we uploaded recordings on Instagram® (version 237.0.0.0.35) and employed a filter named “Black Mask” on each recording. Finally, with a custom Matlab script, we ensured that each pixel within the virtual mask area was black for the entire video duration.

We extracted the audio of each video, which was set to mono, down-sampled to 44100 Hz, and set to 32-bit sample definition using Audacity® (version 2.4.2, https://www.audacityteam.org/). The resultant audios were imported in Matlab (version: R2019b) and equalized to a target Root Mean Square. As a result of a pilot experiment conducted on three participants, we selected the RMS of one story as a target value for all of them (RMS value = 0.03). A separate audio stream of multi-talker babble noise (five voices: two females) was also created. Each babble talker’s voice was recorded in a soundproof room (like the target stimuli) while reading unrelated passages from the fiction book *La Strada* (McCarthy, 2006/2014). These single recordings were equalized to the same mean RMS and then superimposed to obtain the multi-talker babble noise. The multi-talker babble noise was then added to the target speech stories. The first 5 s of the babble were muted (i.e., set to zero), and then the volume linearly increased to generate 5 s of fade-in. This allowed participants to clearly identify the target stories to follow. For the EEG data analysis, we removed the first 11 seconds of each recording to exclude the reading of the title story, the 5 s part in which the babble was muted, and the following 5 seconds in which the babble was fading in. Different segments of multi-talker babble were randomly selected and added to the target stories to avoid adaptation to a specific babble noise. Each target story with babble noise lasted approximately 3 minutes. After piloting, we chose an SNR level of 13.97 dB. iMovie was used to re-pair each preprocessed audio with the video file of the same story and save the obtained final audio-video files in “.mp4” format.

### 2.3 Task and Experimental Procedure

#### 2.3.1 Control to verify participant’s Lip Reading ability with silenced videos

We performed a behavioral experiment to assess the ability of our participants to understand speech content when looking at the videos of the no-mask, virtual mask, and real mask conditions in the absence of sound. For each subject, we presented sixty silent video recordings of the speaker pronouncing single words extracted from our stories. For each participant, twenty video recordings out of sixty were randomly selected for each condition (i.e., no-mask, virtual mask, and real mask). In each trial, a black screen was shown for two seconds, followed by the video recording of a single word presented twice, interleaved by a black screen lasting 2 s. At the end of each trial, participants were required to verbally identify the word they perceived from the video previously shown, always in silence. They were not provided with a list of potential responses; instead, they had to articulate what they comprehended from the silent video presentation. Once the participant performed the behavioral test, the EEG experiment started.

This control experiment was performed to verify whether participants could identify words in the virtual mask and real mask conditions (in the absence of sound) by extracting information from the movements of the mask covering the lips.

#### 2.3.2 EEG Experimental Session

After the behavioral experiment, we started the EEG experiment; each participant attended to audio-visual stories, that is, videos of a speaker talking across no-mask, virtual mask, and real mask conditions. Participants listened to 3 stories per condition randomly presented (see Figure 1). Each story could be presented to each participant only once and could be associated with either no-mask, virtual mask, or real mask conditions, pseudo-randomically. Participants were instructed to listen carefully and maintain fixation at the center of the screen where the speaker’s face was displayed. To help them, a small white fixation cross appeared at the center of the screen before each video. As soon as the experimenter pressed the spacebar, the fixation cross disappeared, and the video started showing the face of the speaker looking at the participant. None of the participants reported knowing the stories.

**Figure 1.**
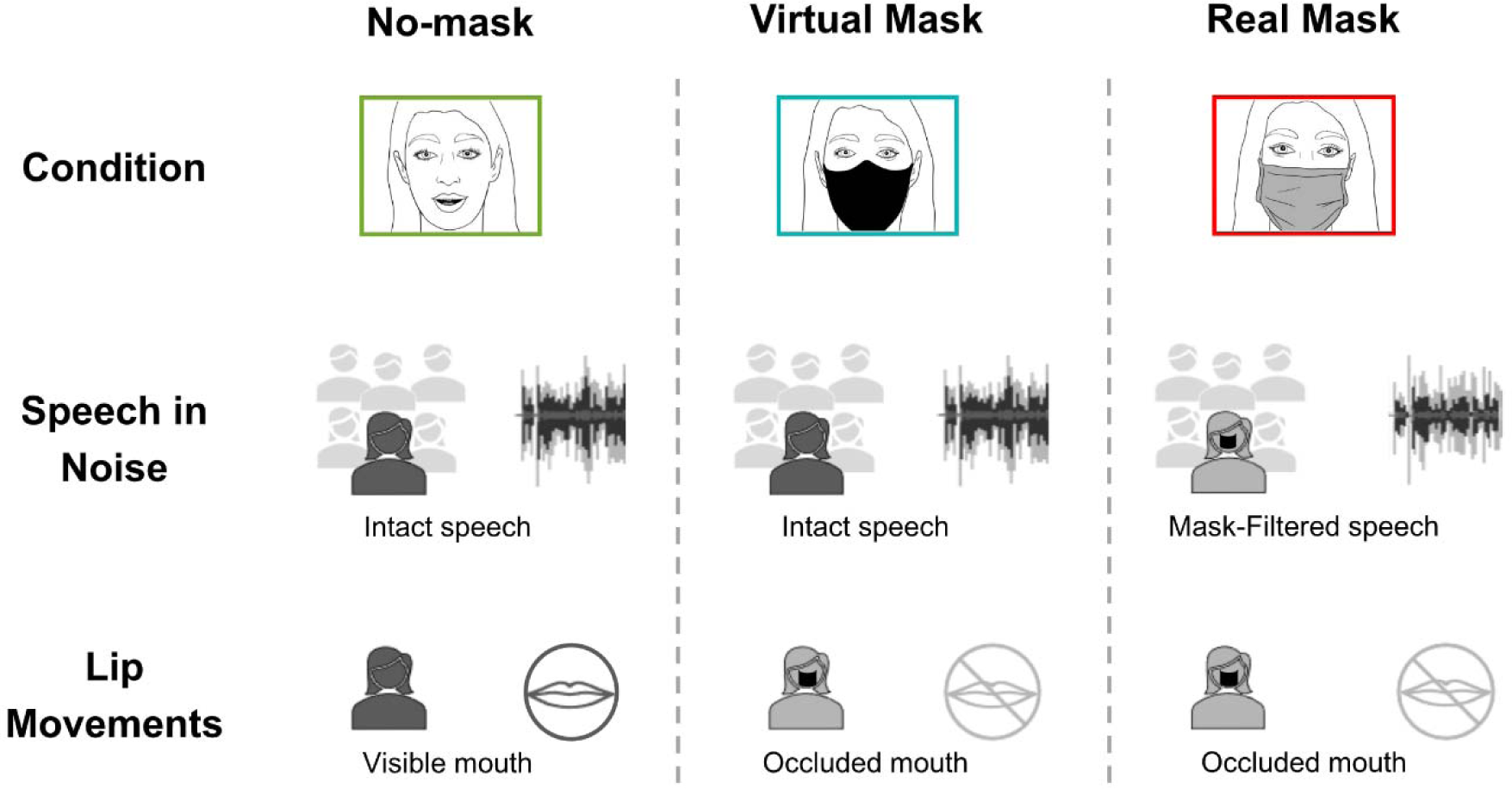
Schematic representation of the three conditions used in the EEG experiment. Participants were attending to continuous audio-visual speech embedded in babble noise (multiple talkers in the background) to promote multisensory integration. In the no-mask condition, visual and acoustic information was fully available to the listener. In the virtual mask condition, the acoustic information was preserved (i.e., identical to no-mask), but a digital face mask hid the lips. In the real mask condition, the acoustic information was degraded by the physical filter constituted by a real-surgical face mask, and the lips were occluded.

Participants were required to answer thirteen questions at the end of each story. The first question was (a) how difficult it was to follow the story on a range from 1 to 7 (1 = not difficult at all, 7 = extremely difficult); (b) six questions concerning the content of the stories to evaluate the retrieval performance, i.e., subject’s accuracy in performing the task. We employed a four alternative forced-choice (4AFC) task. Each question had a correct answer, a wrong answer semantically related to the correct one, a wrong answer phonologically related to the correct one, and an incongruent answer; the order of presentation of the four answers was randomized. (c) A confidence rating followed each question about the content (“*How confident are you about your previous answer?*” they had to answer on a range from 1 to 7, where 1= not confident at all; 7 = extremely confident).

The experiment was performed in a sound-attenuated chamber at the IMT School for Advanced Studies Lucca. Speech stimuli were presented to participants using Psychopy® software (PsychoPy3, v2020.1.3). The sound was delivered using a single front-facing loudspeaker (Bose Companion® Series III multimedia speaker system, country, USA) placed behind the computer screen where the video was displayed. The participant sat approximately 60 cm distant from the screen. Stimuli were delivered at ∼80 dB, measured in front of the loudspeaker (Meterk MK09 Sound Level Meter).

## 3 Analysis

### 3.1 EEG Recording

Participants were asked to remain relaxed and avoid unnecessary movements during the recording session to limit muscle-related artifacts. Blinking was permitted whenever they wanted. The EEG data were recorded continuously during the entire experimental session, using a Brain Products system (BrainVision system ActiCHampPlus) with elastic caps with 32 active electrodes at a sampling rate of 500 Hz (Easy cap Standard 32Ch actiCAP snap). Electrode impedances were kept below the threshold of 30 kΩ. The experiment lasted approximately 1.5 hours per participant, including instructions, EEG Net application, EEG recordings, and breaks when needed by the participants. The alignment of stimulus timing and EEG marker were measured with the AV device (EGI).

### 3.2 EEG Preprocessing

We analyzed only EEG segments in which stories were presented to participants. These segments were concatenated and preprocessed offline with the EEGLab toolbox, Version 14.1.2 (Delorme and Makeig, 2004), and a validated preprocessing pipeline (Stropahl et al., 2018; Bottari et al., 2020). Continuous EEG data were low-pass filtered (cut-off at 40 Hz, Hanning filter order filter 50), downsampled to 250 Hz to reduce computational time, and high-pass filtered (cut-off at 1 Hz, Hanning filter order 500). Then, we segmented the data into consecutive 1-second epochs. The joint probability algorithm removed all the noisy epochs (threshold = 3 SD; Delorme et al., 2007). These filtered and cleaned data were submitted to Independent Component Analysis (ICA, using an EEGLab function based on the extended Infomax (Bell, 1995; Lee, 1999; Jung et al., 2000a, 2000b), and resultant ICA weights were applied to the continuous *raw* data (Stropahl et al., 2018; Bottari et al., 2020). Subsequently, we used CORRMAP, a semi-automatic ICA clustering tool, to identify and remove the components associated with stereotypical artifacts such as blinks or eye movements (Viola et al., 2009). CORRMAP works with a correlation of ICA inverse weights that finds independent components (ICs) similar to a user-defined template. The similarity between the template and each IC is verified by a correlation procedure identifying components exceeding a threshold (r=0.95; Viola et al., 2009). We also inspected the selected ICs using the ICLabel toolbox provided in EEGLAB. Across participants, the mean number of ICs removed by COORMAP was 1.8, SD = 0.4.

Afterward, the cleaned data were low-pass filtered (40 Hz filter order 50), downsampled to 250 Hz, and high-pass filtered (0.1 Hz, Hanning filter order 5000). Noisy channels were detected by applying the automatic bad channel detection algorithm of EEGLab (correlation threshold = 0.9). Deleted channels were interpolated with spherical interpolation (mean channels interpolated across participants = 1.76 ± SD: 1.58). Data were re-referenced to an average reference and band-pass filtered between 2-8 Hz (for 2 Hz cut-off order filter 250, for 8 Hz cut-off order filter 126; see Bednaya et al., 2022; Legendre et al., 2019; Mirkovic et al., 2015; O’Sullivan et al., 2014). Preprocessed EEG data about each story was then epoched. Epochs lasted 3 minutes and started when both target speech and babble noise were present and at stable volume (see stimuli section). Finally, epochs were downsampled to 64 Hz and re-segmented into trials of 1 minute, obtaining 9 trials for each condition and participant. Trials were necessary for the cross-validation procedure; all data were scaled to the max during the evaluation of the regularization parameter (Crosse et al., 2016).

### 3.3 Extraction of Acoustic and Visual regressors

We extracted two regressors related to the visual and the auditory features from our audio-visual stimuli: (i) the lip movements from the video and (ii) the sound envelope from speech (see Supplementary Materials for additional regressors extracted as control features: the Motion and the Clean Audio. Each regressor has been extracted and analyzed separately for each condition.

#### 3.3.1 Lip movements

Lip movements were tracked using DeepLabCut (Mathis et al., 2018). We tracked twelve points: the oral commissures (two) and five parts of the lower and upper lips. For the upper lip, we considered the Philtrum’s movements, the cupid’s bows (two), and the mid-points between the peaks of the cupid’s bows and the lateral commissures. For the lower lip, we considered the upper lips’ mirror counterparts along the lower vermillion border (See Supplementary Materials and Figure S1). For each point, we extracted the X and Y positions (in pixels) for each frame and calculated the mouth’s area. In this way, we measured the change in lip movements over time. The lip movements were downsampled to 64 Hz and epoched according to the EEG Preprocessing (3 minutes for each story, including only the portions of the recordings where the audio of both target speech and babble noise were present and at stable volume). Epochs were then cut into 1-minute long trials, obtaining three trials for each story, i.e., nine trials in total, in each condition. Signals were then normalized. Normalization was carried out for neural and stimulus data, following Crosse et al. (2021). This approach was applied to normalize stimulus features to prevent variations in feature scales that could impact the magnitude of feature weights

#### 3.3.2 Sound envelope

To extract the sound envelope, we imported the audio streams in MATLAB. As for the lip movements, we epoched each story and cut files into 1-minute-long trials (N=9 trials in total). Then, the audio envelope was extracted for each trial taking the absolute value of the Hilbert transform of the original stories (i.e., the envelope was estimated using the clean stories without multi-talker babble noise; Kortelainen and Vayrynen, 2015), applying a low-pass filter with a third order Butterworth filter (with 8 Hz cut-off; *filtfilt* MATLAB function), and downsampling to 64 Hz to match the EEG data resolution (O’Sullivan et al., 2015; Mirkovic et al., 2015). Finally, we normalized the envelope by dividing each value by its maximum value to optimize the cross-validation necessary to estimate the regularization parameter (Crosse et al., 2016; Bednaya et al., 2022).

### 3.4 Stimulus Reconstruction

To estimate the reconstruction of different speech features (lip movements and sound envelope) from the neural data, we employed a backward model using the mTRF Toolbox (Multivariate Temporal Response Function, see Supplementary Materials; Crosse et al., 2016). This linear model reconstructs the speech feature of interest from the EEG activity recorded in all electrodes; importantly, the sensors’ activity is weighted according to their informativeness (Crosse et al., 2016; Holdgraf et al., 2017).

By applying this model, we wanted to estimate associations between the EEG signals and the stimuli speech features, including typical predictions occurring before the unfolding of the stimulus in the case of continuous language (de Lange et al., 2018; Heilbron et al., 2022).

In this model, the time lags equal to zero represent the exact synchrony between brain activity and speech features. Thus, at negative or positive lags, the model estimates the association between the brain activity ahead or in late with respect to speech features.

To train the model, a leave-one-out cross-validation was performed to find our optimal regularization parameter, *λ* (for more details, see Supplementary Materials: “Regularization Parameter Estimation”). Once we tuned the model parameter, *λ*, the model was tested on the same data set (Crosse et al., 2016). We trained the decoder with the chosen lambda with leave-one-out cross-validation, keeping apart one trial at each iteration, and then we performed the mean of the models obtained at each iteration. Afterward, we tested the averaged model on each left-out trial, and we computed the reconstructed speech feature (lip movements or sound envelope). Then, we correlated the reconstructed speech feature to the original one; the result of this Pearson correlation is the reconstruction performance (Decoding r).

This procedure was performed for each decoding model calculated across time lags in a range between -115 and +575 ms, with a sliding time window of 45 ms and an overlap of 30 ms with adjacent time lags (Mirkovich et al., 2016). The decoding was calculated in each time window, obtaining an estimate of the averaged r value across participants at the center of each time window. For example, in the case of the first time window, we had an estimate of the r value between -115 and -70 ms. After that, we selected three 50-ms windows of interest according to the reconstruction performance peaks on which we performed the decoding model and then transformed in the forward direction to become neurophysiologically interpretable and to be shown in the results images. Indeed, it is impossible to neurophysiologically interpret the resulting decoder weights topography (Haufe et al., 2014). Transformed decoder weights have a more straightforward interpretation; their value and sign are directly related to the strength of the signal at different channels (Crosse et al., 2021).

#### 3.4.2 Estimation of the Null Effect (null decoding)

We computed a null decoding model for each participant and condition to test whether the decoder’s performance was above chance. To this end, we permuted the order of 1-minute trials of the original speech feature (lip movements or sound envelope) to obtain mismatched speech feature and EEG response pairs on which the decoding was fitted to estimate the null decoding (mTRFpermute function with 100 iterations; Crosse et al., 2016, 2021). Then, the correlation coefficients of all these decoding models, computed across iterations, were averaged to obtain the ‘null decoding.’ This procedure was done separately for each participant.

### 3.5 Statistical Analysis

#### 3.5.1 Report of the p-values

In the behavioral results (both behavioral control and behavioral test within EEG experiment), having performed single tests, all p-values, both Bonferroni corrected (p_corr_) or uncorrected p-values (p), have been reported as exact p-values.

Instead, in the decoding results, since we performed multiple tests, all the statistically significant p-values, both FDR corrected (p_FDR_) or uncorrected p-values (p), have been reported as <0.05. The non-statistically significant p-values have been reported as >0.05.

For significant effects, the effect size (d’Cohen; Cohen, 1988; Cousineau, 2021) and the confidence intervals (CIs) have been reported to evaluate the magnitude of the observed effects. The CIs have been computed by performing the bootstrap method with N=1000 bootstrap replicas.

#### 3.5.2 Behavioral control: Lip Reading ability of silenced videos

The behavioral test on lip-reading allowed us to evaluate the participants’ ability to comprehend silent speech across the different conditions (no-mask, virtual mask, and real mask). A non-parametric Friedman test (one factor, three levels) was employed to control the fact that the occlusion of the lips reduced lip reading ability. Multiple comparison post-hoc tests were performed with Bonferroni correction.

#### 3.5.3 Behavioral test within the EEG Experiment

A Friedman test was used as the primary statistic to test the behavioral outcomes of the EEG experiment. Post-hoc tests were performed by applying the multiple comparison tests with Bonferroni correction; the corresponding p-values were reported as p_corr_ when statistically significant.

We evaluated (1) the participant’s *accuracy* in answering content retrieval questions and the relative (2) committed *errors* (which will be described in detail subsequently), (3) the *confidence* of the participants when answering the content questions, (4) the *perceived difficulty* by participants when listening to speech embedded in babble noise.

As mentioned, we wanted to explore further the type of errors committed by participants when answering the content questions. Participants had to select among four alternative forced choices (4AFC), and the answers were differentiated into (1) *correct* answers, (2) answers *semantically* related to the correct one, (3) answers *phonologically* related, and (4) *incongruent* answers.

#### 3.5.4 Statistical Analysis of the Decoding Outcomes

First, we assessed the existence of neural entrainment to the lip movements and sound envelope at the group level. To this end, we compared the reconstruction performance at the group level of each of our regressors against their null decoding performance, i.e., decoding vs. null decoding within each condition. Analyses have been conducted by transforming the correlation coefficient values (r; of both the real decoding, indicating the correlation between reconstructed and original stimulus, and the null decoding, showing the correlation between reconstructed and the mismatched stimulus) of each condition through Fisher’s transformation, obtaining z transformed values. The z values of each condition were then compared with one-tailed paired t-tests at each time lag (-115 +575 ms). P-values were corrected across all time lags (N = 47) using the False Discovery Rate (FDR) procedure to control for multiple comparisons (Benjamini, 1995).

Second, we contrasted the decoding z values across conditions. Here, we directly compared the reconstruction performance values between conditions with a series of one-tailed paired t-tests across all time lags in which we measured reliable reconstruction performance (where the real decoding was significantly higher than the null decoding). We computed each t-test under the specific hypotheses that reconstruction performance would have the following directionality: no-mask (full cues) > virtual mask (visual obstacle) > real mask (visual and acoustic obstacle). P-values were corrected for multiple comparisons (time lags) with the FDR procedure (Benjamini, 1995).

#### 3.5.5 Analysis of the Correlation between Neural and Behavioural Data

After establishing whether neural tracking of the sound envelope and the behavioral measures varied across conditions, we were interested in analyzing potential correlations between neural tracking and behavioral outcomes.

To achieve this, only in case significant differences emerge between conditions, we conducted a correlation analysis between the reconstruction performance and the behavioral data. These correlations were computed at each time lag. Subsequently, we examined both uncorrected and corrected p-values, the latter applying a correction for multiple comparisons across time lags (FDR procedure proposed by Benjamini in 1995).

## 4. Results

### 4.1 Control on Lip Reading ability of silenced videos

As expected, the Friedman test revealed that participants were severely impaired at recognizing words when the lips were hidden (χ^2^_Friedman_ (2)=51.37, p<0.001; N=30 participants, mean age=27.7 years; 15 females). Post-hoc tests (Bonferroni corrected) confirmed a significant difference between no-mask and the two face mask conditions (no-mask vs. virtual mask conditions p_corr_< 0.001; d’Cohen=1.86, CI [1.35 2.39]); no-mask vs. real mask conditions p_corr_< 0.001; d’Cohen=1.83, CI [1.35 2.44]); no significant difference emerged between virtual and real mask conditions (p_corr_=1, see Supplementary Materials Figure S2).

### 4.2 Behavioral test results of the EEG experiment

Results on participants’ *accuracy* in answering content questions did not reveal significant differences across conditions, suggesting that even in mask conditions, participants are able to attend to the auditory input efficiently. Indeed, individuals’ ability to process and retrieve speech content embedded in multi-talker babble noise did not change across conditions (^2^ (2)=5.3, p=0.07). However, the mask condition modulated the kind of errors participants committed when answering the content questions.

Participants had four alternative forced choices (4AFC), the correct answer, and three wrong answers. The wrong answers could be phonologically or semantically related to the correct answer or completely incorrect. A statistically significant difference emerged in *phonological* errors (χ^2^_Friedman_ (2)=10.14, p=0.006); the post hoc test (Bonferroni corrected) revealed a selective difference between no-mask and real mask conditions (p_corr_=0.004; d’Cohen= 0.75, CI [0.39 1.18]; see Figure 2b). This result suggested that the presence of a surgical face mask covering the lips and filtering the auditory information increased errors in which wrong answers were phonologically associated with correct responses (no significant effects for the comparisons across the other conditions; no-mask vs. virtual mask: p=0.33; virtual mask vs. real mask: p=0.33;). Conversely, neither *semantic* (χ^2^_Friedman_ (2)=1.77, p=0.41) nor *incongruent* errors were modulated by the type of condition (χ^2^_Friedman_ (2)=1.15, p=0.56; see Figure 2b).

**Figure 2.**
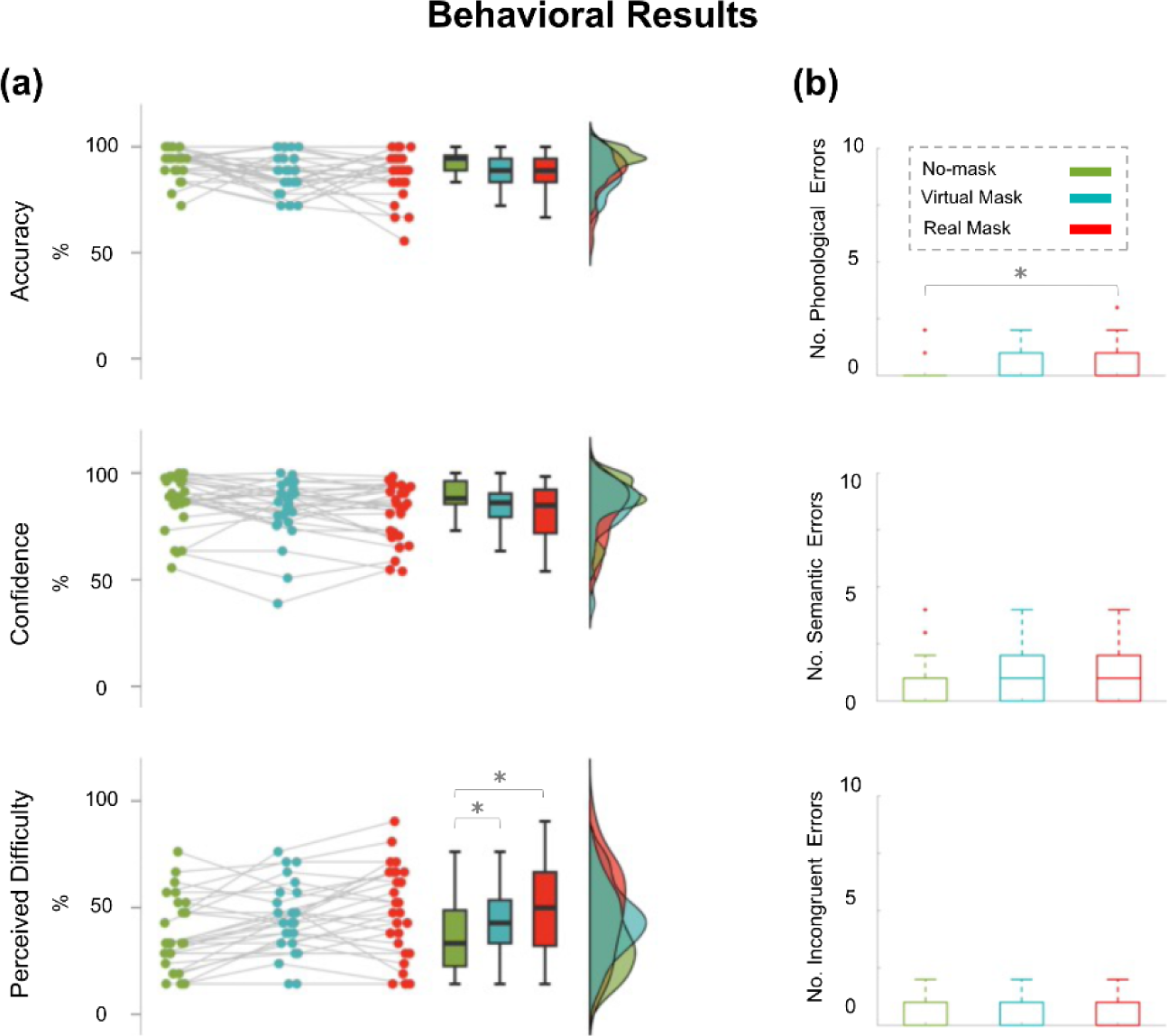
(a) Results of the three behavioral measures acquired during the EEG experiment (*accuracy*, *confidence*, and *perceived difficulty*) shown across conditions. Listening to a person wearing a face mask (virtual or real) increased participants’ perceived difficulty. (b) Type of errors performed in the content questions: on the top, the phonological, middle, and semantic, and on the bottom, the incongruent. In the real mask condition, the number of errors committed increased significantly when the answers were phonologically associated with the correct responses.

Participants’ *confidence* when answering each content question was not affected by the mask condition (χ^2^_Friedman_ (2)=5.39, p=0.06).

Finally, results on the perceived *difficulty* highlighted a significant main effect of the condition (χ^2^_Friedman_ (2)=17.11, p<0.001). Post-hoc tests (Bonferroni corrected) revealed a statistically significant difference between the no-mask and the virtual mask condition (no-mask vs. virtual mask p_corr_=0.006 d’Cohen=0.44, CI [0.16 0.71]) and between the no-mask and the real mask condition (no-mask vs. real mask p_corr_<0.001; d’Cohen= 0.57, CI [0.31 0.85]); but no difference between virtual and real mask conditions (p_corr_=1). Results indicated that the presence of a mask hiding the lips, both virtual or real, substantially increased perceived difficulty following speech embedded in multi-talker babble noise.

### Decoding Model Results

#### 4.3.1 Lip Movements Decoding Results

First, we evaluated the decoding of lip movements in the no-mask and virtual mask conditions (the virtual mask videos were created by applying a virtual mask on the actor’s face recorded in the no-mask condition videos, and thus, the lip movements, despite being invisible, had precisely the same kinematic). We obtained successful lip movement reconstruction only in no-mask condition. Indeed, reconstruction performances exceeded the null decoding at time lags from 50 to 575 ms (p_FDR_<0.05; d’Cohen=1.03, CI [0.71 1.43]; see Figure 3). Conversely, we obtained no significant results in the virtual mask condition (p_FDR_>0.05), suggesting that we found no evidence for participants reconstructing lip movements from acoustic information. Additionally, we investigated the decoding of a control visual model in the no-mask condition (the Motion Energy; Nishimoto et al., 2011) to verify if it was possible to decode the information related to the displacement of the mouth area. In this case, we found that it was possible to decode general motion information, but the lip movements were indeed more informative (see Supplementary Materials Figure S3).

**Figure 3.**
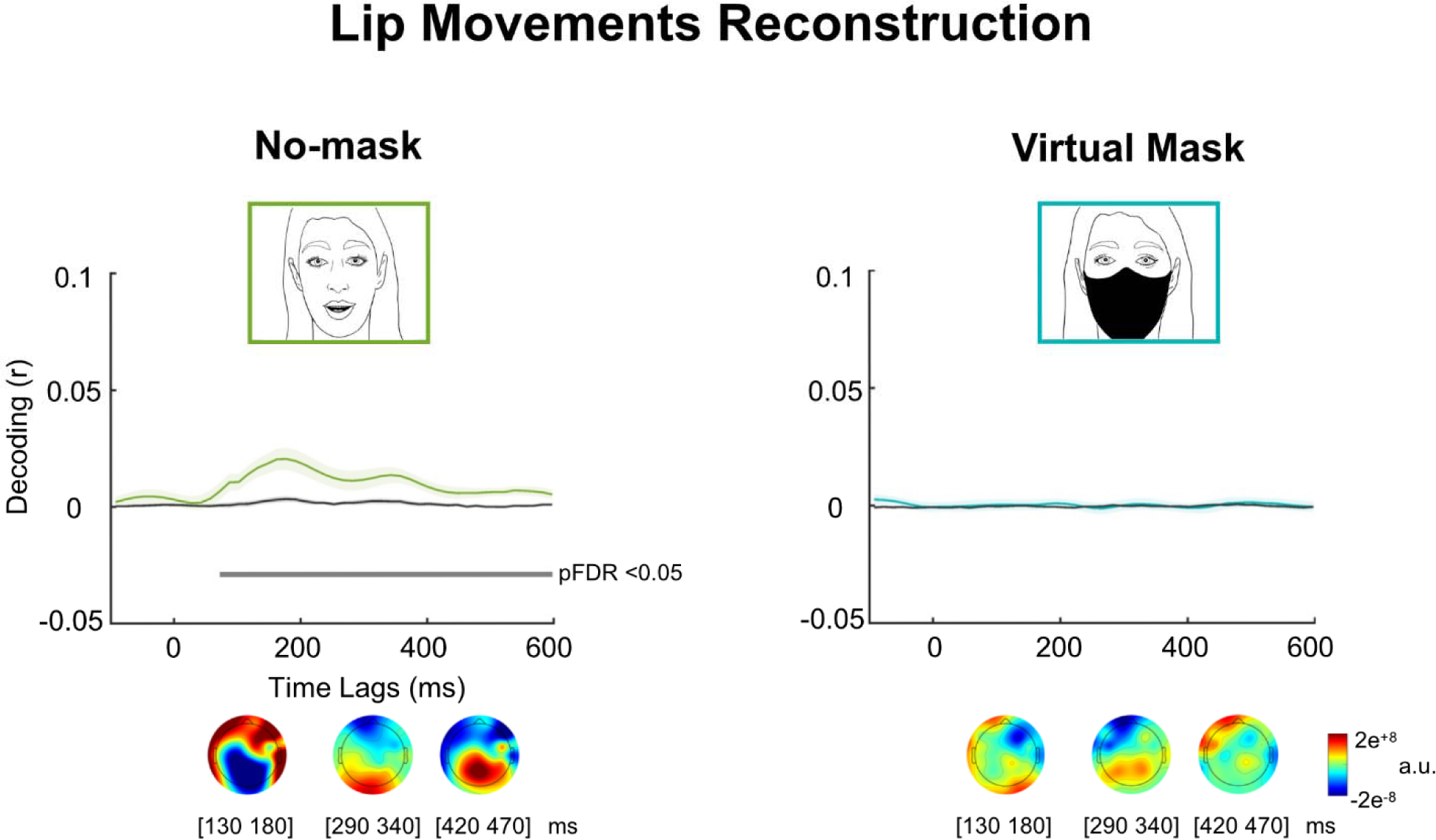
Neural decoding of the lip movements in no-mask (left) and virtual mask conditions (right). The correlation coefficients of the lip movements represent the reconstruction performance - Decoding (r), on the y-axis - calculated as the correlation between the reconstructed and original lip movements at each time lag; the corresponding null decoding is depicted in grey. The x-axis comprises time lags between -115 and +575 ms. The continuous line represents the group mean, and the shaded areas represent the SE. Grey horizontal lines under the null decoding highlight the statistically significant time lags in which the reconstruction performance exceeded the null decoding of the envelope (FDR corrected statistics). Only for the no-mask condition we observed a reliable reconstruction of the lip movements from the EEG data. R values are plotted at the center of each time lag from ∼72 and at last ∼597 ms). Under each plot, forward topographies for each condition are depicted at representative 50 ms time windows.

#### 4.3.2 Sound Envelope Decoding Results

For each condition, we estimated the neural tracking of the sound envelope by comparing the reconstruction performance of auditory stimulus features against their corresponding null reconstruction, also defined as null decoding. The reconstruction performance values exceeded the null decoding in each condition (no-mask condition: -115 to 410 ms, p_FDR_<0.05; d’Cohen=2.58, CI [1.48 3.34]; virtual mask condition: -115 to 425 ms, p_FDR_<0.05; d’Cohen=2.54, CI [1.91 3.32]; real mask condition: -115 ms up to 320 ms, p_FDR_<0.05; d’Cohen=2.01, CI [1.47 2.52]). Results clearly provided evidence of successful decoding of sound envelope irrespective of whether the mouth was visible and/or speech was filtered by a face mask (see Figure 4a).

**Figure 4.**
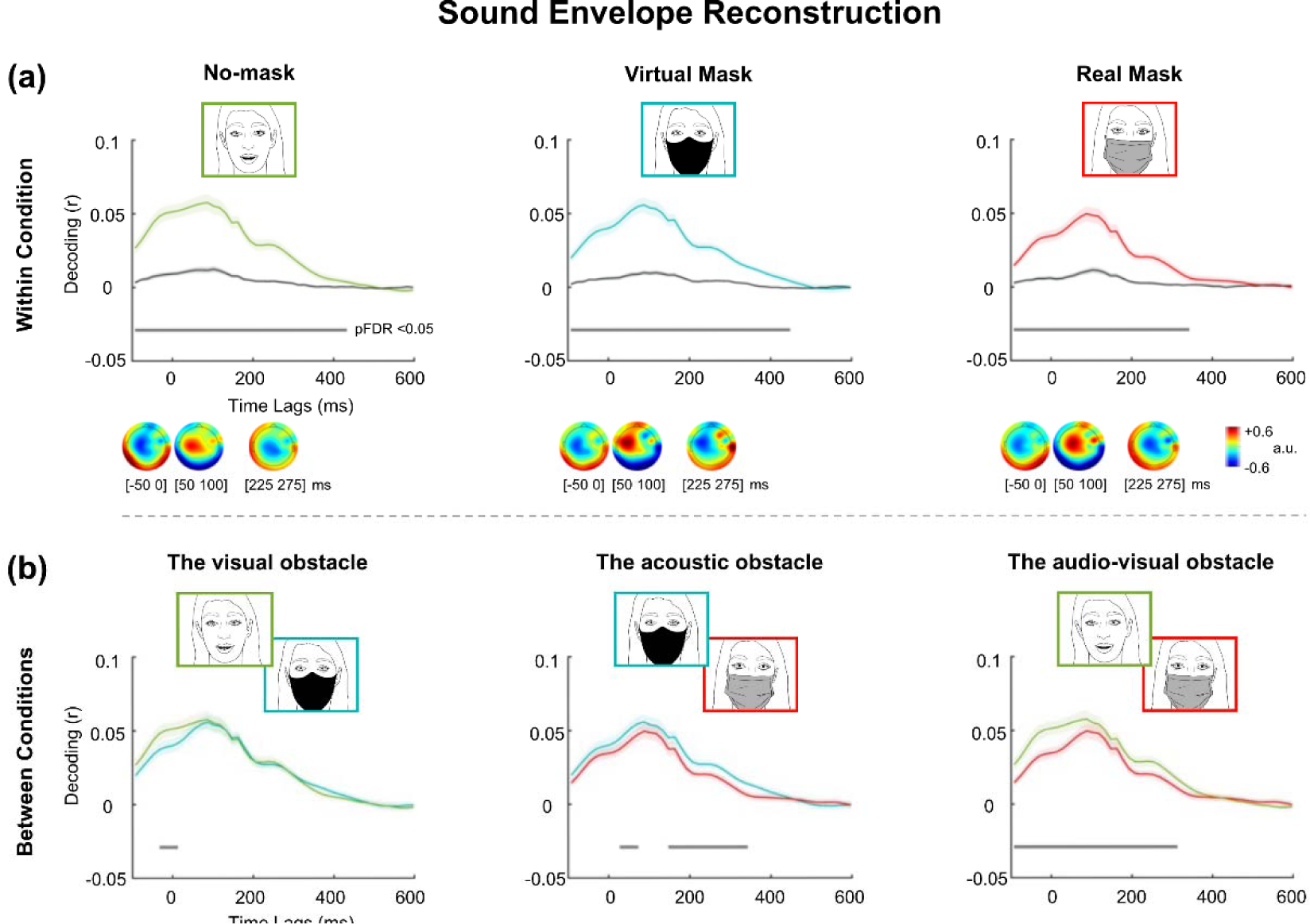
(a) Sound envelope reconstruction in no-mask, virtual, and real mask conditions. The correlation coefficients represent the reconstruction performances - Decoding (r), on the y-axis - calculated as the correlation between the reconstructed and original sound envelope at each time lag; the corresponding null decoding performances are depicted in grey. The x-axis comprises time lags between -115 and +575 ms. The continuous line represents the group mean, and the shaded areas represent the SE. Grey horizontal lines under the null decoding highlight the statistically significant time lags in which the reconstruction performance exceeded the null decoding of the envelope (FDR corrected statistics). In each condition, we measured reliabl reconstruction performance. R values are plotted at the center of each time lag (no-mask vs. null decoding between ∼ -92 and ∼432 ms, virtual mask vs. null decoding between ∼ -92 and ∼447 ms, and real mask vs. null decoding between ∼ -92 and ∼342 ms). Below each plot, forward projected topographies of decoding models at representative 50 ms time windows. (b) The *visual obstacle* effect was investigated by comparing the no-mask vs. the virtual mask condition (left, r values plotted at center between ∼ -32 and ∼12 ms), the *acoustic obstacle* effect was observed by analyzing the virtual mask vs. the real mask condition (center, r values plotted at center between ∼27 and ∼72 ms and from ∼147 and ∼342 ms), and the combined *audio-visual obstacle* effect was obtained by contrasting the no-mask vs. the real mask condition (right, r values plotted at center between ∼92 and ∼312 ms). For all comparisons, the statistically significant time lags, that is, the time points in which the neural tracking differed between conditions, are represented by a grey line (FDR corrected statistics).

Having demonstrated reliable sound envelope reconstructions, we investigated the impact of auditory and visual obstacles provided by face masks. To this aim, we contrasted the decoding accuracy across conditions. First, we tested the no-mask vs. virtual face mask condition to estimate the specific impact of covering the lips on the neural tracking of the sound envelope (the acoustic information was identical across these two conditions), that is, the impact of the *visual obstacle*. The decoding of the sound envelope in the no-mask condition exceeded the virtual mask condition at the earliest time lags, between -55 to -10 ms (p_FDR_<0.05; d’Cohen=0.43, CI [0.08 0.79]; see Figure 4b), suggesting that available lip movements contributed to the neural tracking by anticipating the sound envelope tracking.

We then specifically estimated the *acoustic obstacle* determined by the face mask. To do so, we compared the reconstruction performance of the sound envelope between the virtual mask and the real mask conditions. In both conditions, the lips were not visible due to the presence of a face mask, but only in the real mask condition was the auditory input filtered by the surgical mask, as the speech was recorded while the speaker was wearing it. Sound envelope reconstruction was higher in the virtual mask as compared to the real mask at several time lags (between 5 to 50 ms and between 125 and 320 ms; p_FDR_<0.05; d’Cohen=0.49, CI [0.21 0.81]; see Figure 4b), suggesting that the physical barrier constituted by the surgical mask hampered the auditory neural tracking at later processing stages as compared to the *visual obstacle* (mouth occlusion).

Finally, we contrasted the reconstruction performance of the sound envelope between no-mask and real mask conditions to estimate the combined impact of lips occlusion and acoustic filter. Results revealed a protracted dampening of reconstruction performance between -115 and 290 ms (p_FDR_<0.05; d’Cohen=0.57, CI [0.31 0.91]; see Figure 4b). This effect revealed the combined *audio-visual obstacle* generated by listening to a speaker wearing a surgical face mask.

Having measured the different obstacles on neural tracking generated by face masks, we further investigated the importance of the shared information between lip movements and sound envelope in the absence of any obstacle. To this aim, we created a new regressor, the clean audio, in which we regressed out all the common information between the lip movements and the sound envelope (this was possible only in the no-mask condition). By comparing the decoding of cleaned audio regressor and the standard sound envelope in the no-mask condition, we selectively described the auditory neural tracking at the net of information provided by audio-visual correspondences (between lip movements and sound envelope). A significant difference emerged, with a long-lasting decrease in reconstruction performance -115 to 380 ms (see Supplementary Materials Figure S4), highlighting the role of the common information between lip movements and sound envelope dynamics for the neural tracking of speech. The decoding drop occurred at time lags substantially overlapping with the combined *audio-visual obstacle* generated by listening to a speaker wearing a surgical face mask.

### 4.4 Correlation between Neural and Behavioural Data

At the behavioral level, face masks significantly affected only the perceived difficulty in attending speech. Here, we aimed to explore whether a relationship existed between this behavioral measure and neural tracking. In each condition, we analyzed the correlation between the reconstruction performance of the sound envelope at each time lag (at the single participant level, for each time lag, decoding -r- were transformed into z values with Fisher transform), and the related perceived difficulty. An association emerged between the two mask conditions. In the virtual mask condition, correlation coefficients were statistically significant between 200 and 245 ms (p<0.05); in the real mask condition, between 170 and 230 ms (p_FDR_<0.05; see Figure 5).

**Figure 5.**
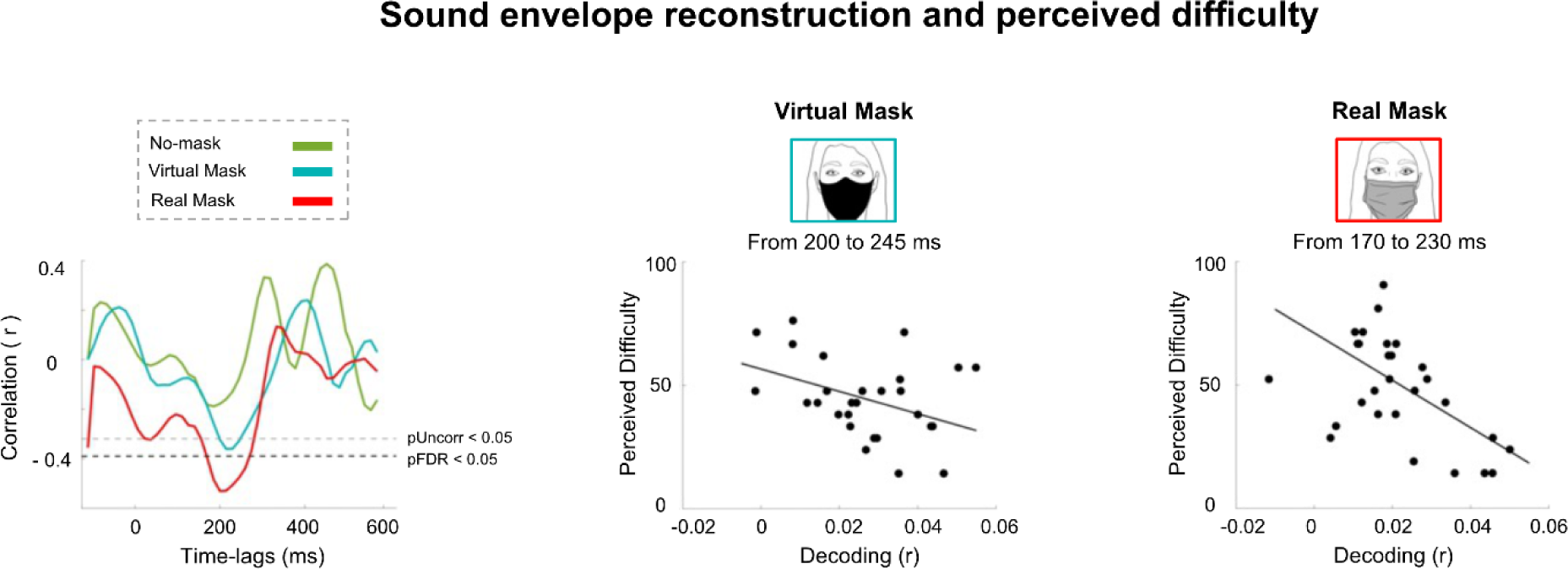
Correlations at each time lag between the neural tracking of the sound envelope with the perceived difficulty in each condition (left panel). A negative correlation between the neural tracking and the perceived difficulty was found in the two mask conditions (virtual and real mask) at time lags ∼200 ms.

## 5. Discussion

In the present work, we dissected auditory and visual obstacles generated by face masks on face-to-face communication. Face masks increased perceived difficulty in attending the speech, and when the speaker wore a real surgical mask, there were greater phonemic errors when listeners were asked to report communication content. As expected, we observed that covering the mouth area prevented the neural tracking of lips. Conversely, sound envelope reconstruction was possible in all conditions. Yet, when contrasting the decoding accuracy of sound envelope across conditions, we observed a progressive dampening of reconstruction performance depending on whether we measured the visual, the acoustic, or the combined filtering effect. Finally, we found that the difficulty of listening when the speaker wears a face mask was mirrored by the costs measured at the neural level.

### 5.1 Behavioral effects of face masks on speech processing

No significant changes in *accuracy* in answering speech content questions emerged across the different conditions, suggesting that the SNR was high enough to allow efficient speech processing in all tested contexts. This outcome aligns with previous studies showing that face masks do not substantially affect speech comprehension (Toscano and Toscano, 2021; Haider et al., 2022). Similarly, participants’ *confidence* was unaffected when answering content questions. Noteworthy, we observed a difference in the kind of errors committed by participants when answering; results suggested that *phonological* errors increased when listening to a speaker wearing a surgical face mask, that is when both visual and auditory obstacles were combined. Previous findings outlined that when listening to a speaker, we process parallel visual and acoustic information, which support speech comprehension (Chandrasekaran et al., 2009). Seeing the articulatory movements of the speaker’s face sustains intelligibility (Holler et al., 2019), especially in challenging listening conditions (Moradi et al., 2013; Puschmann et al., 2019). Moreover, speech processing is hampered when the acoustic information is degraded, such as by the presence of face masks (Corey et al., 2020; Choi et al., 2022), which are known to dampen frequencies above 1-2 kHz (with different effects depending on the material used for the mask; Corey et al., 2020).

Moreover, we found a progressive increase of *perceived difficulty* across conditions, depending on whether only the lip information was absent (no-mask vs. virtual mask) and when both visual and auditory inputs were altered (no-mask vs. real mask). This was coherent with previous studies highlighting the impact of face masks on metacognitive levels of communication (Giovanelli et al., 2021). In recent years, various studies used face masks as a model to investigate audio-visual speech integration (Brown et al., 2021; Rahne et al., 2021; Thibodeau et al., 2021; Toscano et al., 2021; Choi et al., 2022). Results revealed that there is no difference in understanding a speaker wearing a face mask or just listening to the voice, once more stressing the lack of audio-visual benefit on speech comprehension if lips are occluded (Giovanelli et al., 2021).

### 5.2 The neural tracking of lip movements and sound envelope

First, we analyzed the neural tracking of the visual information in the no-mask and virtual mask conditions. Lip movements could be successfully reconstructed only in the no-mask condition, between 50 and 575 ms, highlighting activity at occipital sensors. Conversely, we measured neural tracking of the sound envelope for all conditions at central sensors. In the no-mask and virtual mask conditions, we had robust and prolonged sound envelope tracking up to -115 and 425 ms. In the real mask condition, it was from -115 to 320 ms, instead.

Time lags at which decoding of lip movements was successful were delayed compared to the sound envelope reconstruction dynamic. This agrees with results showing that lip movement processing is more protracted than a sound envelope (Holler et al., 2019). A recent MEG study showed that the brain can reconstruct acoustic information from silent lip movements (Hauswald et al., 2018). If it were possible to decode the lip movements even in the virtual mask condition, it would have indicated that the brain reconstructs the unseen lip movements from heard speech, but we found no such evidence (but see Bourguignon et al., 2020).

When listening to speech in noise, like in a multi-talker situation, paying attention to the target sound of speech and filtering out unattended voices implies a greater effort for the listener (O’Sullivan et al., 2014; Ahmed et al., 2023). Especially in noise, the availability of both acoustic and visual speech signals is fundamental for efficient tracking and comprehension (Crosse et al., 2016b). Indeed, looking at the lips greatly supports speech tracking and intelligibility (Park et al., 2016; Giordano et al., 2017). A recent MEG study showed that surgical face masks negatively affect listeners’ neural tracking of acoustic features (i.e., sound envelope and spectral features; Haider et al., 2022). The authors interpreted these results as the consequence of the missing visual input and the subsequent impossibility of integrating acoustic and visual information. Nonetheless, to fully understand the mechanisms underlying this effect, it is crucial to disentangle the impact of face masks’ acoustic and visual filters. The main novelty of this study lies in the clear separation of the contributions provided by audio and visual speech features for the neural tracking of speech. Based on previous findings (Bourguignon et al., 2020; Tan et al., 2022; Haider et al., 2022), we expected a decreasing reconstruction performance of speech envelope along a continuum spanning from no-mask, virtual, and real masks.

### 5.3 Masking the lips impairs early reconstruction of the sound envelope

We analyzed the specific *visual obstacle* on the sound envelope reconstruction, that is, how the visual filter of the face mask undermined speech processing. To this aim, we compared the no-mask vs. virtual mask conditions and measured a drop in the reconstruction performance of the envelope at time lags between -55 and -10 ms.

Acoustic and visual information are correlated in continuous speech, with a peak in frequency bands corresponding to the syllabic rate between 4 and 8 Hz (e.g., Park et al., 2016). Evidence of the influence of visual cues on acoustic processing is given by the cross-modal phase modulation of the auditory entrainment by visual speech signals. Power and colleagues (2012) evaluated the neural tracking of speech signals and analyzed the role of congruent visual speech cues on auditory entrainment. The phase of auditory tracking was altered in the presence of congruent visual stimulation. Visual speech also represents a critical complement to auditory speech signals by providing information on temporal markers of the upcoming acoustic stimulus and the relative content. Coherent visual cues support the neural tracking of the auditory cortex, enhancing speech processing (Peelle, Sommers, 2015). This phenomenon is an example of multimodal facilitation of speech processing (Drijvers and Holler, 2023). When available, visual cues decrease the listener’s cognitive load and enhance the precision in predicting speech (Peelle, Sommers, 2015). Moreover, visual cues convey information about the position of the speaker’s articulators that become fundamental when listening to speech in noise (Peelle, Sommers, 2015), such as in the present experiment. The brain can efficiently integrate audiovisual signals of speech despite lip movement onsets preceding the sound onsets with a range of about 100 to 300 ms (Chandrasekaran et al., 2009). Our findings are coherent with this; here, we observed a precocious detrimental effect of the lack of visual information over sound envelope tracking (no-mask vs. virtual mask) at the earliest time lags due to the role of visual cues on early stages of speech processing.

### 5.4 Uncovering the effect of the acoustic filter

By contrasting the virtual vs. real mask condition, we assessed the impact of the degraded auditory information on the sound envelope tracking, namely the *acoustic obstacle*. In this comparison, we selectively evaluated the sound filtering effect of a real face mask since the mouth area was covered in both conditions. Given that listening to speech in noisy environments demands focused attention on target speech and suppress distractors (Mesgarani et al., 2012; Kim et al., 2021), we predicted more favorable neural tracking in the virtual mask condition compared to the real mask condition, as the latter represented a more compromised scenario. Coherently, we found that the envelope tracking in the virtual condition exceeded the one in the real mask condition at several time lags, from 5 to 50 ms and 125 to 320 ms. The main difference between the speech envelope reconstruction occurred around 200 ms, a typical time scale of neural tracking associated with auditory selective attention (e.g., O’Sullivan et al., 2014).

Protective face masks have been of fundamental importance during the pandemic of COVID-19; despite that, consequent difficulties in face-to-face communication have been underlined, especially for listeners with hearing impairment (Corey et al., 2020; Homans, Vroegop, 2021). Previous behavioral studies highlighted the impact of face masks in filtering acoustic information, leading to decreased comprehension both in normal hearing and hearing-impaired participants (Choi et al., 2022; Homans Vroegop, 2021; Moon et al., 2022). Also, face masks challenge auditory attention, intelligibility, and memory recall, especially when listening to speech in noise (Rimmele, 2015; Smiljanic et al., 2021). According to the mask’s fabric, the detrimental effect on comprehension and subjective listening effort can vary (Brown et al., 2021; Thibodeau et al., 2021). Overall, the present results confirmed and expanded such evidence by providing their possible neural correlates (Toscano and Toscano, 2021; Brown et al., 2021).

### 5.5 Real surgical masks: acoustic and visual obstacles combined

The contrast between the no-mask vs. real mask condition allowed us to evaluate the effect of the combined *acoustic-visual obstacles*. Here, we found that the decoding of the sound envelope in the no-mask condition exceeded the real mask within time lags between -115 and 290 ms. As described earlier, face masks affect speech at multiple levels: by covering the mouth area (Corey et al., 2020) and preventing lip reading, by hampering the acoustic propagation of the voice (Thibodeau et al., 2021), and affecting the subjective perceived difficulty and metacognitive performance monitoring (Giovanelli et al., 2021). Coherently, we observed that the *acoustic-visual obstacle* led to the greatest and most prolonged dampening of speech neural tracking. To further dissect the role of audio-visual information on the tracking of the sound envelope, we also developed a predictor (clean audio) from which we regressed out shared information between the lip movements and the sound envelope. This allowed estimating neural tracking without the influence of audio-visual correspondences. Coherently with the previously described combined *audio-visual obstacle*, the decoding drop associated with the removal of shared audio-visual information occurred at similar times, as in the case of listening to a speaker wearing a real surgical face mask (see Supplementary Material).

### 5.6 Link between Neural and Behavioral data

We explored potential associations between the perceived difficulty in attending speech and the decoding of the sound envelope as a function of the different face mask conditions. Notably, negative correlations between behavioral and neural measures were observed in the virtual mask condition from 200 to 245 ms and in the real mask condition between 170 and 230 ms time lags. Conversely, no significant results were found in the no-mask condition. Results demonstrated a link between the progressive decline of neural tracking depending on the perceptive load induced by the mask conditions and the increase in the perceived difficulty in listening to speech. The association between neural activity and metacognition emerged at time lags at around 200 ms, typically indicative of auditory attention processes (e.g., O’Sullivan et al., 2014). While a positive relationship between speech intelligibility and the tracking of acoustic features is well known (see Ding and Simon 2013), these results are consistent with recent evidence illustrating a connection between neural tracking of sound envelope and perceived difficulty (Reisinger et al., 2023).

## 6. Conclusions

In the current study, we developed a naturalistic paradigm based on continuous speech to investigate how the brain synchronizes with speech signals in detrimental listening conditions (face masks and multi-talker noise). Results allowed disentangling between the acoustic and visual impact of face masks on speech processing. These obstacles had impacts at partially distinct processing stages. Findings substantiated the role of audio-visual congruent information in speech processing and showed an increased load on metacognition in the case audio-visual processing of speech is hindered. These results highlight the impact of face masks on social communication and have established objective indicators that can serve as a reference for developing and validating new materials used to construct these devices, aiming to reduce perceptual obstacles and metacognitive load. These findings have the potential to offer objective tools to optimize future choices, develop better protective devices, and - ultimately - contain the communication problems linked to the use of masks by improving the quality of life.

## Acknowledgments

The authors want to thank Dr. Chiara Valzogher for her contribution to the first draft of the manuscript.

