## Supplementary Material for "The impact of face masks on face-to-face neural tracking of speech: auditory and visual obstacles"

**Supplementary Materials**

1. **Stimulus Reconstruction: The Backward Model**

As previously described, to evaluate the reconstruction performance of different speech features from the recorded EEG data (the lip movements and the sound envelope), we employed the mTRF Toolbox (Crosse et al., 2016). We used the decoding model, also called the “backward model”, to predict selected speech features from the neural data separately. In Figure S1, the model is illustrated.

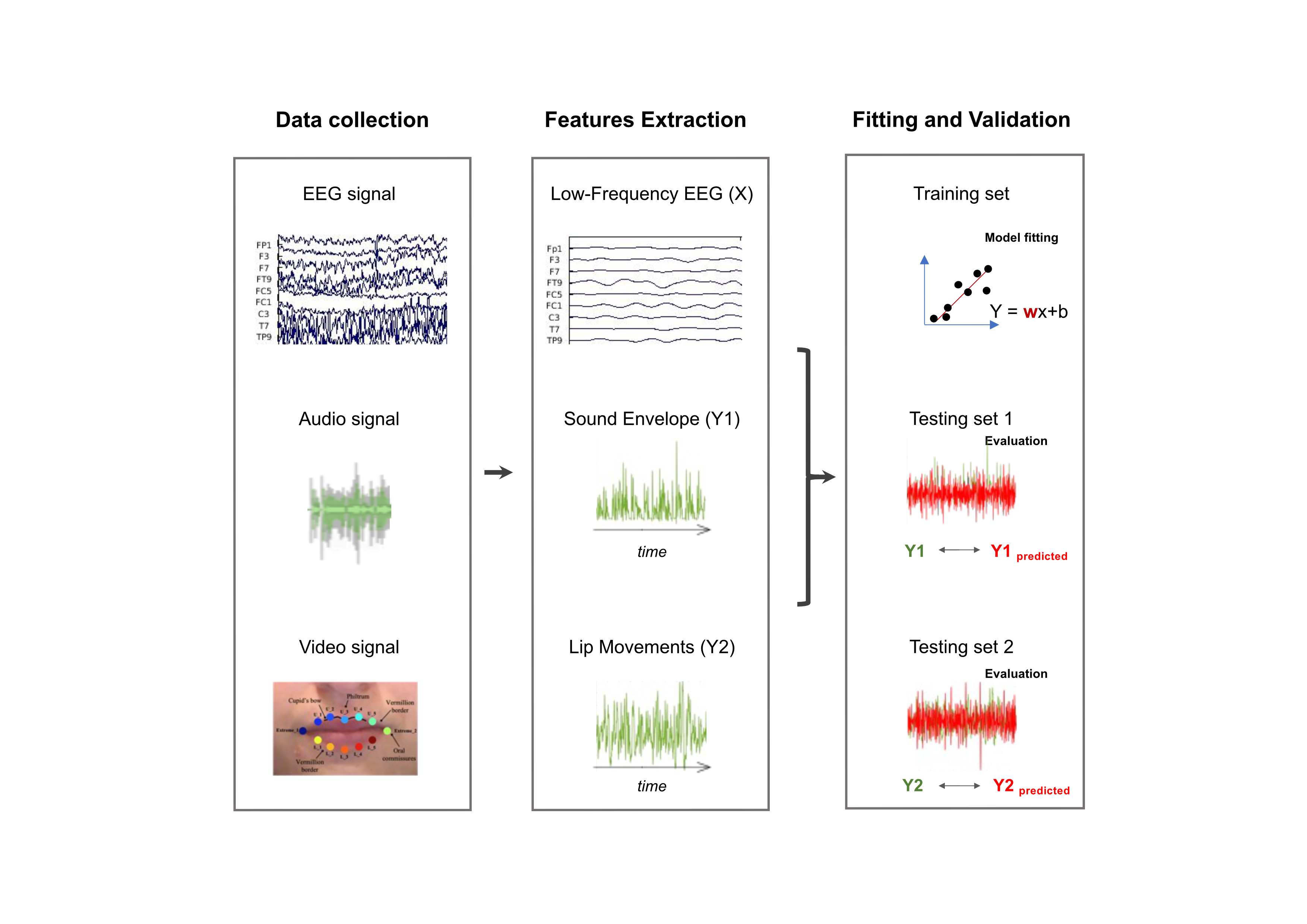

Figure S1. Description of stimulus reconstruction approach. We decoded speech features (sound envelope and lip movements) from EEG data to quantify the linear relationship between each speech feature and the neural activity. We pre-processed EEG data to obtain the cleaned EEG signal, the audio signal to extract the sound envelope, and the video of the speaker to extract lip movements (note, other features were also extracted by these naturalistic stimuli and are described in the next paragraph). Each model was separately trained to reconstruct the corresponding stimulus feature (i.e., sound envelope and lip movements) from the neural data. The model was tested by evaluating the degree of correlation between reconstructed and original speech features.

2. **Regularization Parameter Estimation**

To avoid overfitting (Crosse et al., 2021), we applied the best possible ridge parameter ($\lambda$) over the optimal time lag to obtain the best decoder able to predict stimulus features. For each participant, we tuned the regularization parameter using the leave-one-out cross-validation procedure, using a pre-defined series of possible values (i.e., λ = 10^-3^, 10^-2^, 10^-1^, … ,10^3^, 10^4^, … , 10^8^, 10^9^, 10^10^) to compute the model for time lags from -100 to 600 ms (Crosse et al., 2021). Lambda was chosen based on the mean squared error (MSE) value. The procedure returns the mean squared errors MSE that was used as a criterion to choose the final $\lambda$ value; indeed, the best lambda returned the lowest MSE on the testing data (here, it was $\lambda$ = 10^4^ for most subjects). This value was kept constant across participants and conditions, allowing us to generalize results at the group level.

3. **Control on Lip Reading ability of silenced videos: Results**

To test the participants’ lip-reading ability, we showed them sixty silenced videos in our three conditions: no-mask, virtual mask, and real mask. In each video, the face of the speaker was presented while telling a single word in one of the three conditions; each video was presented twice, with a brief pause (2 sec) between the first and the second presentation. In Figure S2, the results are reported; a statistically significant difference emerged between the no-mask and the two-mask conditions. From this result, we can conclude that participants were highly affected by the presence of a face mask (virtual or real) when recognizing words without sound, determining a significant drop in performance with respect to the no-mask condition.

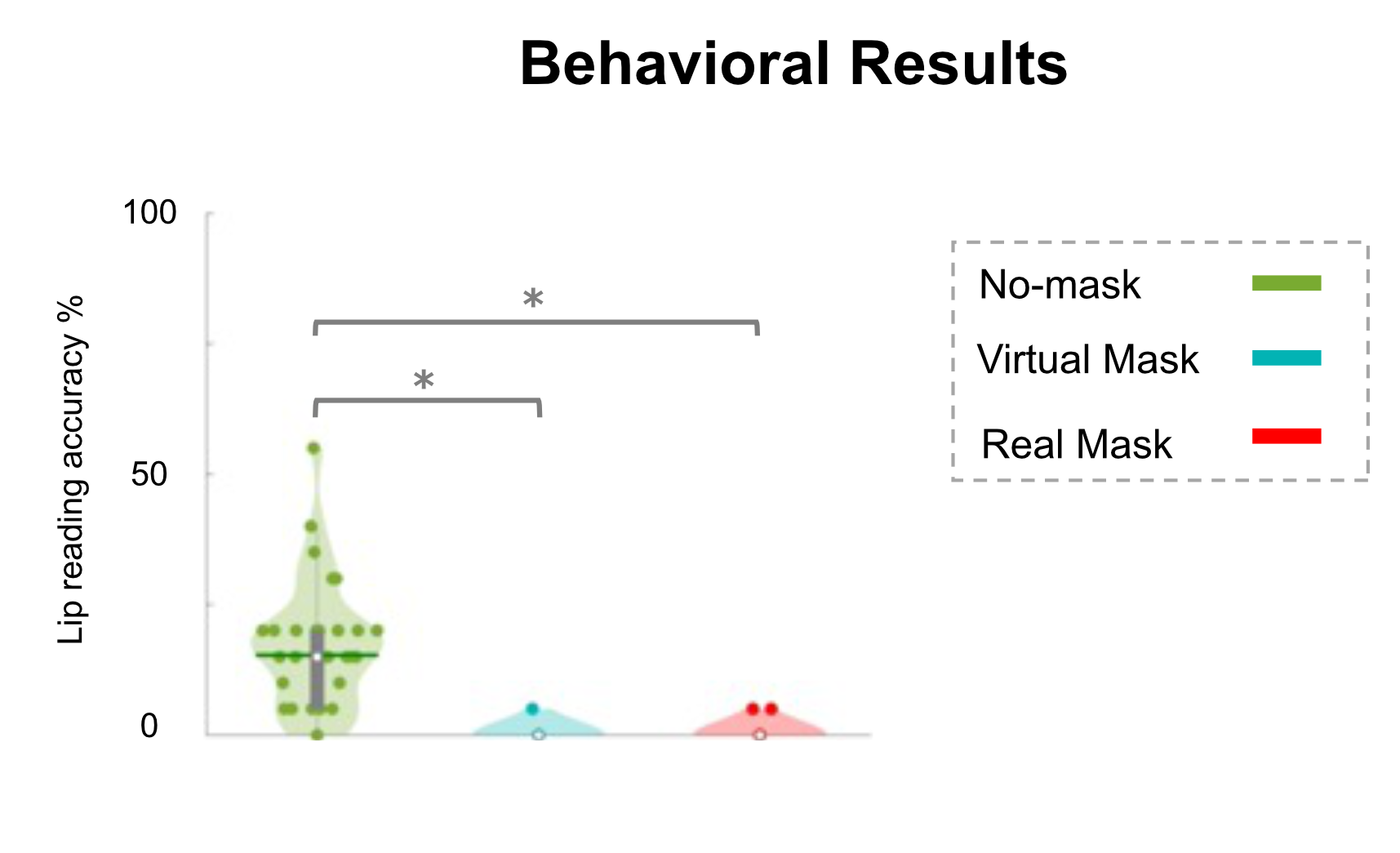

Figure S2. Results of the control on lip-reading ability in the three conditions; results are shown in accuracy %.

**3. Audio-Visual Control regressors: the Motion energy** **and the Clean Audio**

In addition to the lip movements and sound envelope, we estimated the reconstruction of two other regressors as controls: (1) the motion energy to assess the decoding related to the general movements occurring in the mouth area, and not just the lip movements, (2) and the clean audio to evaluate the tracking of the sound envelope at the net of the information related to the visual cue.

**3.1 Report of the p-values**

All the statistically significant p-values, both Bonferroni corrected (p_corr_), FDR corrected (p_FDR_) have been reported as “<0.05”. Instead, the non-statistically significant p-values have been reported as “>0.05”.

**3.2 Pre-processing of the Motion energy** **and the Clean Audio regressors**

After pre-processing the lip movements and sound envelope, we assessed the regressor for the motion energy regressor, we reduced the size of our videos for computational purposes; then, we cropped our recordings to show only the mouth area (size of the cropped videos: columns = 480, rows= 270). Then, to compute the motion energy, we applied the procedure illustrated in Nishimoto et al. (2011) and the relative customized scripts available on GitHub: <https://github.com/gallantlab/motion_energy_matlab>. In our pipeline, we obtained a matrix with the RGB values of each pixel for each frame in time. Then, we converted the pixels to greyscale and applied the Gabor wavelet processing (we used customized parameters suitable for our stimuli and purposes; we reduced the number of features, the SR value was set to 30 Hz, and the frequencies of interest were defined between 2 and 8 Hz). Then, we computed the principal component analysis (PCA) and selected the first component as our motion regressor. After that, we downsampled our data to 64 Hz and cut the first 11 seconds of the regressor; also, the outliers were identified and substituted. This regressor consisted of the averaged motion energy across all pixels in the mouth area.

To pre-process the acoustic information in isolation, i.e., the clean audio. To do so, we first computed the average rank of the two regressors (lip movements and sound envelope) for further processing. Then, we defined a sliding integration time window of 468.7 ms (corresponding to 30 frames at 64Hz) that has been calculated referring to the work of Stevenson and Wallace (2013). The authors described the temporal binding window, TBW (i.e., the temporal interval for integrating stimuli temporally related) for different sets of stimuli, including audio-visual speech. Then, we computed the correlation between the ranked lip movements and the sound envelope based on the sliding integration window. After that, we assessed the regressor of the clean audio by multiplying the sound envelope (AudioEnv) by one minus the absolute value of the correlation (Rho) between the sound envelope and the lip movements.

$$\mathrm{CleanAudio}=(\mathrm{AudioEnv}(1:end-30))\cdot(1-\mid\mathrm{Rh}o(1:end)\mid)'$$

In this way, we removed the shared information between lip movements and sound envelope from the regressor dynamic with the aim of evaluating how the brain tracked the acoustic cue without the beneficial effect of seeing the lips.

**3.3 Decoding Model Results of the Motion Energy regressor**

Then, we assessed the motion energy neural tracking of the mouth area in the no-mask condition; indeed, it was the only condition where participants had complete visual information. We contrasted the reconstruction performance of the motion regressor vs. its null decoding. In this comparison, we obtained statistically significant results from 20 to 365 ms, and from 530 to 545 ms (p_FDR_<0.05; d’Cohen=1.03, CI [0.57 1.49]; see Figure S4, the panel on the left). This outcome shed light on decoding visual features with different regressors from the mouth area. Once we had verified that it was possible to decode the motion cue, we wanted to investigate if there were statistically significant differences between the stimulus reconstruction of the visual input using different visual regressors: the motion energy and the lip movements (see Figure S4, panel on the right). To do so, we compared the reconstruction performances obtained with these two regressors, finding statistically significant differences from 95 to 200 ms with p_FDR_<0.05; d’Cohen=0.78, CI [0.28 1.27]. To conclude, we found that the neural tracking of the lip movements was greater than the one of the motion cue at different time lags.

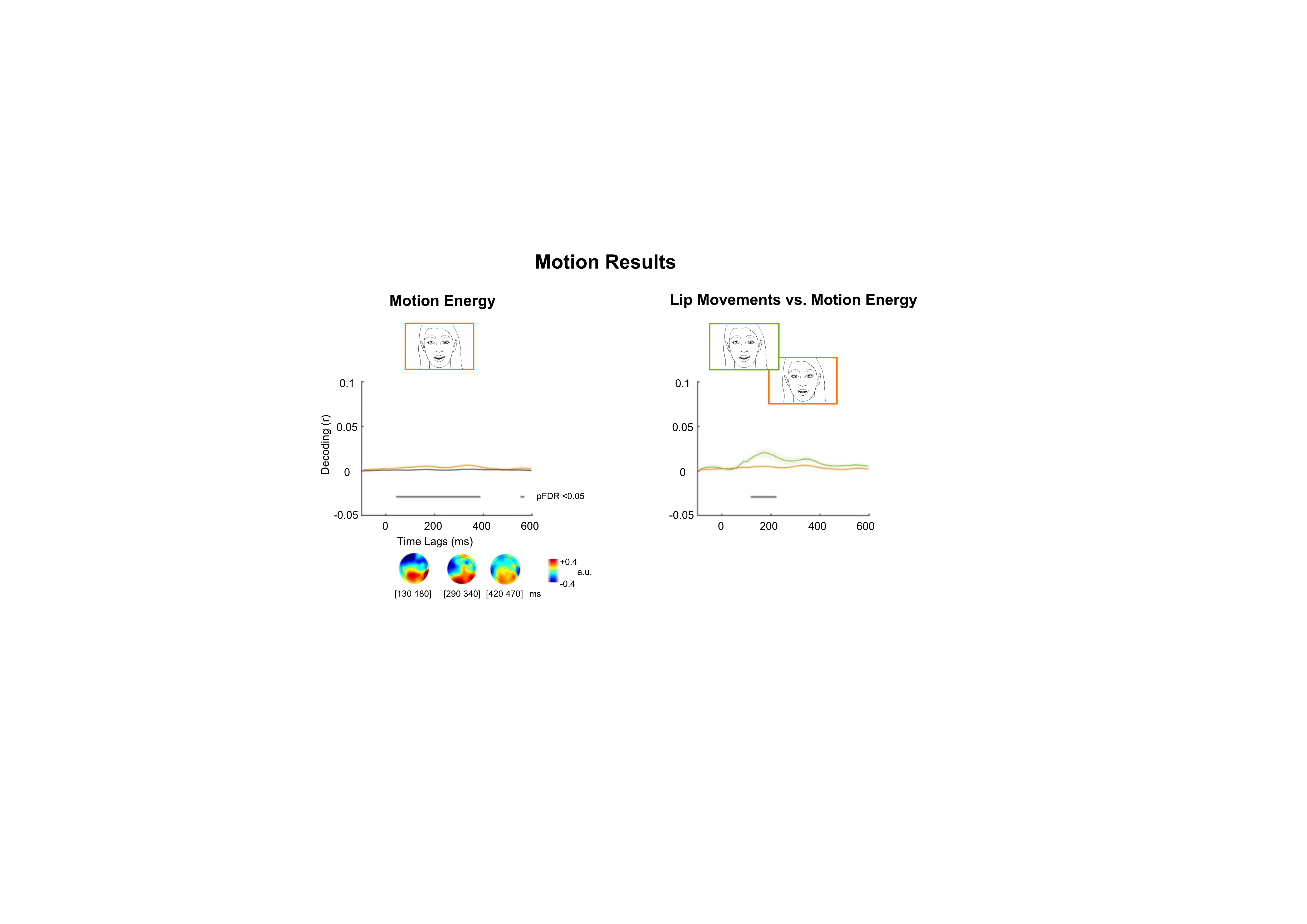

Figure S3. Neural decoding of the motion regressor in no-mask on the left, the comparison between the motion and the lip movements regressor in no-mask condition on the right. The correlation coefficients of the motion and of the lip movements are represented as reconstruction performances - Decoding (r), on the y-axis - calculated as the correlation between the reconstructed and original feature at each time lag; the corresponding null reconstruction performances are depicted in grey. The x-axis comprises time lags between -115 and +575 ms. In the first plot on the left, we evaluated the reliable reconstruction for the motion regressor; the grey horizontal line under the null decoding highlights the statistically significant time lags in which the reconstruction performance of the motion exceeded the null reconstruction of the envelope (FDR corrected statistics), r values are plotted at the center of each time lag from ~42 and ~387 ms and from ~552 and at last ~567. Below, forward projected topographies for the motion depicted at representative 50 ms time windows. On the plot on the right, instead, we contrasted the reconstruction performance of the motion with the reconstruction of the lip movements in no-mask condition; here, the grey horizontal line represents when the reconstruction performance of the lip movements exceeded the reconstruction of the motion. Also in this case, r values are plotted at the center of each time lag from ~117 and at last ~222 ms.

The motion energy has been applied in the literature to obtain estimates explaining the motion information of the stimuli provided to the participants. For example, Jessen and colleagues (2019) compared the auditory and motion brain responses between infants of 7 months and adults. In that case, the authors computed the motion regressor as the averaged motion across all the pixels of the video recordings presented to the participants. In the current analysis, we showed that the tracking of the motion energy associated with the mouth area was possible. However, we found that lip movements were far more informative than the general motion energy, providing a better reconstruction of the visual input.

**3.4 Decoding Model Results of the Clean Audio regressor**

We analyzed the clean audio envelope in the no-mask condition, where participants could hear the acoustic cue properly and look at the visual one. By contrasting the reconstruction performance of the clean audio (that is, the sound envelope ‘cleaned’ from the shared information between the acoustic and the visual stimulus) vs. its null decoding, we obtained statistically significant results from -115 up to 380 ms (p_FDR_<0.05; d’Cohen=2.28, CI [1.45 2.98]; see Figure S3, panel on the left). This result revealed successful decoding of the pure acoustic information from an audio-visual condition. Also, we were interested in evaluating if there were differences within the no-mask condition between the tracking of the sound envelope (already described in the paper) vs. the tracking of the clean audio (See Figure S3, panel on the right). By comparing the two reconstruction performances, we found a significant difference from -115 to 305 ms, with p_FDR_<0.05; d’Cohen=0.31, CI [0.21 0.45]. The stimulus reconstruction of the sound envelope exceeded the clean audio in no-mask condition, an effect due to the removal of the shared audio-visual information from the decoding model. This outcome was in line with previous literature (Crosse et al., 2015; Park et al., 2016; Bourguignon et al., 2020). In addition, the forward topographies had a similar distribution to one of the sound envelope described in previous analyses, with greater activity in correspondence with the central sensors.

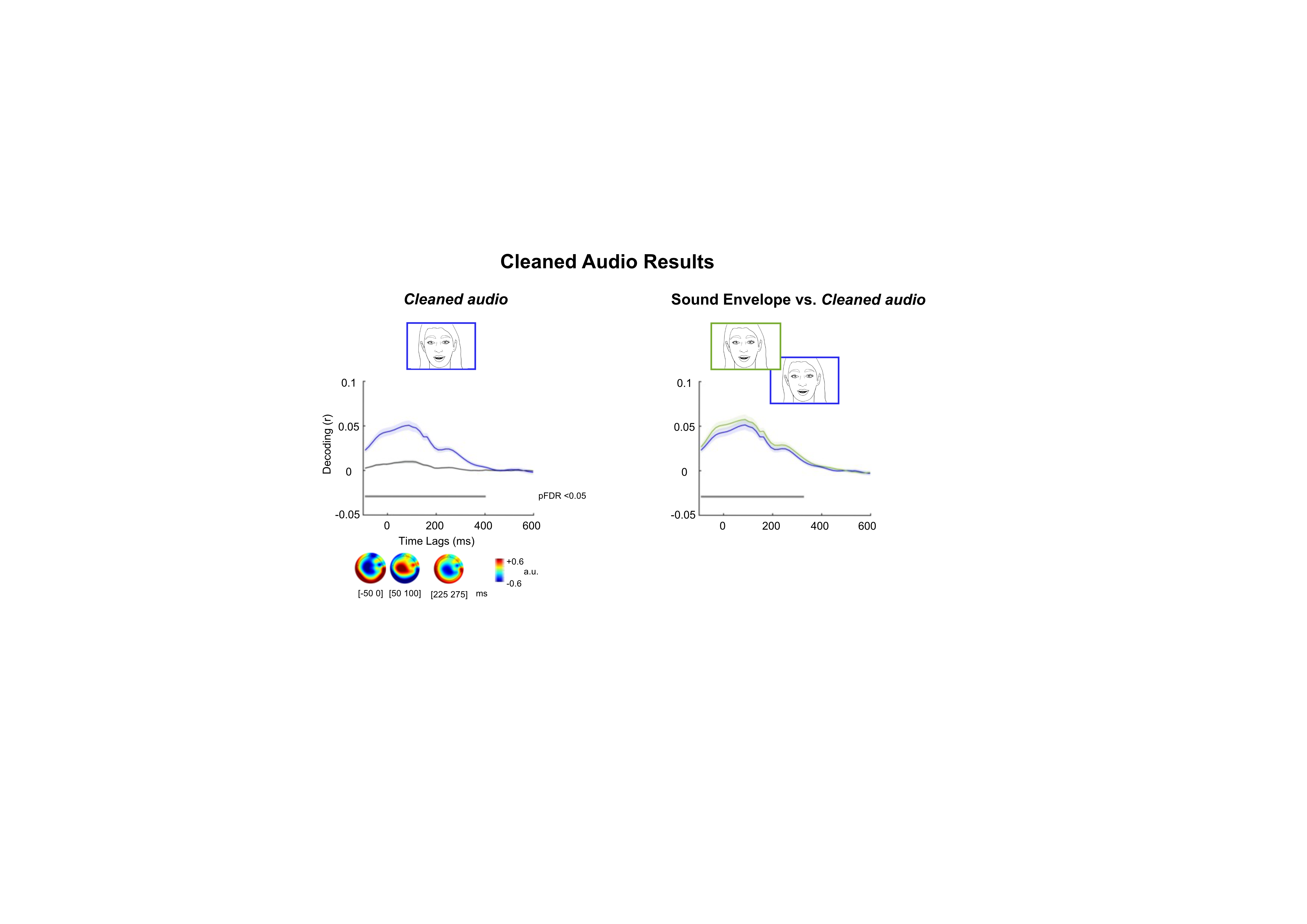

**V1**

**Versione presente nel paper**

**Ho modificato I titoli su suggerimento di Alessandra**

**Next slide vedi quelli vecchi**

Figure S4. Neural decoding of the clean audio regressor in no-mask on the left, the comparison between the sound envelope and the clean audio regressor in no-mask condition on the right. The correlation coefficients of the motion and of the lip movements are represented as reconstruction performances - Decoding (r), on the y-axis - calculated as the correlation between the reconstructed and original feature at each time lag; the corresponding null reconstruction performances are depicted in grey. The x-axis comprises time lags between -115 and +575 ms. The continuous line represents the group mean, and the shaded areas represent the SE. In the first plot on the left, we evaluated the reliable reconstruction for the clean audio regressor; the grey horizontal line under the null decoding highlights the statistically significant time lags in which the reconstruction performance of the clean audio exceeded the null reconstruction of the envelope (FDR corrected statistics), r values are plotted at the center of each time lag from ~92 and at last ~402. Below the plot, forward projected topographies for the clean audio are depicted at representative 50 ms time windows. On the plot on the right, instead, we contrasted the reconstruction performance of the clean audio with the reconstruction of the sound envelope in no-mask condition; here, the grey horizontal line represents when the reconstruction performance of the sound envelope exceeded the reconstruction of the clean audio. Also in this case, r values are plotted at the center of each time lag from ~92 and at last ~327 ms.
